# Modulation of metabolic hormone signaling *via* a circadian hormone and a biogenic amine in *Drosophila melanogaster*

**DOI:** 10.1101/2020.09.25.312967

**Authors:** Jason T. Braco, Cecil J. Saunders, Jonathan M. Nelson, Erik C. Johnson

## Abstract

In insects, Adipokinetic hormone is the primary hormone responsible for the mobilization of stored energy. While a growing body of evidence has solidified AKH’s role in modulating the physiological and behavioral responses to metabolic stress, little is known about the upstream endocrine circuit that directly regulates AKH release. We evaluated the AKH-expressing cell transcriptome to identify potential regulatory elements controlling AKH cell activity, and found that a number of receptors show consistent expression levels, including all known dopamine receptors, dopamine ecdysone receptor (DopEcR), Dopamine 2-like receptor (D2R), Dopamine 1-like receptor 2 (DopR2), DopR, and the Pigment Dispersing Factor (PDFR). We tested the consequences of targeted genetic knockdown and found that RNAi elements targeting each dopamine receptor caused a significant reduction in survival under starvation. In contrast, PDFR knockdown significantly extended lifespan under starvation whereas expression of a tethered PDF in AKH cells resulted in a significantly shorter lifespan during starvation. These manipulations also caused various changes in locomotor activity under starvation. Specifically, there were higher amounts of locomotor activity in dopamine receptor knockdowns, in both replete and starved states. PDFR knockdown resulted in increased locomotion during replete conditions and locomotion levels that were comparable to wild-type during starvation. Expression of a membrane-tethered PDF led to decreased locomotion under baseline and starvation. Next, we used live-cell imaging to evaluate the acute effects of the ligands for these receptors (dopamine, ecdysone, and Pigment Dispersing Factor) on AKH cell activation. Dopamine application led to a transient increase in intracellular calcium in a sugar-dependent manner. Furthermore, we found that co-application of dopamine and ecdysone led to a complete loss of this response, suggesting that these two hormones are acting antagonistically. We also found that PDF application directly led to an increase in cAMP in AKH cells, and that this response was dependent on expression of the PDFR in AKH cells. Together these results suggest a complex circuit in which multiple hormones act on AKH cells to modulate metabolic state.

## Introduction

Throughout the Metazoa, metabolic homeostasis is coordinated by a number of different hormones that coordinate between storage and release of energy. In *Drosophila*, insulin-like peptides (dILPS) and Adipokinetic hormone (AKH), modulate energy storage and release, respectively (Kim and Rulifson 2004; Leopold and Perrimon 2007). The cells that express these hormones represent important points of integration that couple changes in metabolic allocation towards different behaviors and/or physiologies (*e.g*., reproduction or growth). While many studies have focused on the regulation of the dILPS (e.g., Toivonen and Partridge 2009; Birse *et al.* 2011; Kannan and Fridell 2013), there has been comparatively little research identifying elements that regulate the synthesis, release, and signaling of AKH.

Genetic ablation of AKH producing cells (APCs) (Lee *et al.* 2004; Braco *et al.* 2012), blockade of AKH hormone release (Braco *et al.* 2012), or the loss of the AKH or the AKH receptor gene (AKHR) (Galilkova et al., 201, Yu, Huang, Ye, Zhang, Wang, *et al.* 2016), all produce consistent metabolic and behavioral phenotypes. Specifically, these manipulations cause an increase in triglyceride content, and reduce levels of circulating trehalose (Lee *et al.* 2004; Isabel *et al.* 2005). The AKHR, which binds AKH with high affinity, is expressed at high levels in the fat body and genetic knockdown of AKHR in this tissue produces many of the same metabolic phenotypes associated with the loss of the AKH hormone (Grönke *et al.* 2007; Bharucha *et al.* 2008). In animals, a common behavioral response to nutrient depletion is increased locomotion (Scheurink *et al.* 2010; Mistlberger 2011). This may seem counterintuitive as hyperactivity during nutrient limitation would exacerbate energy depletion, however, it is hypothesized that starvation-induced activity is an adaptive trait that facilitates foraging (Johnson 2017). Interestingly, animals lacking AKH show no changes in locomotor activity under starvation conditions (Lee *et al.* 2004). Consequently, AKH mutants have increased lifespan under starvation, presumably caused by the lack of starvation-induced hyperactivity and elevated levels of energy stores (Lee *et al.* 2004).

Despite our growing understanding of AKH signaling, little is known about the factors that govern AKH release. Currently, only two mechanisms are known which detect changes in intracellular energy in this cell lineage. AKH cells express two energy sensors: K^+^_ATP_ channels and AMPK (Kim and Rulifson 2004; Braco *et al.* 2012). These molecules couple to changes in cellular ATP with electrical excitability and AKH release (Stephan *et al.* 2006). While these two molecules are critical factors modulating AKH release, they are unlikely to be the sole mechanisms. Recent work has shown that the loss of AMPK only partially phenocopies the loss of AKH, suggesting alternative or additional mechanisms controlling AKH secretion (Braco *et al.* 2012).

Studies in other insects implicate a number of diverse hormones are modulating AKH release. Crustacean cardioacceleratory peptide (CCAP), tackykinin (TK), and octopamine are positive regulators of AKH release and FLRFamide and FMRFamide act to inhibit AKH release in the locust (Pannabecker and Orchard 1986; Brown and Lea 1988; Veelaert *et al.* 1997; Vullings *et al.* 1998). In general, these experiments measured AKH titer in entire CNS preparations or whole organism in response to hormone application, thus making it impossible to assert whether these hormones are acting directly to alter AKH release or if they are acting through some intermediate (Pannabecker and Orchard 1986; Brown and Lea 1988).

Here, we report the identification of the complement of peptide and amine receptors expressed in the AKH cell lineage employing single cell RNA sequencing methods. We further assessed the functional contributions of the PDF receptor, and all known dopamine receptors with specific regards to modulation of AKH cell physiology. Genetic knockdown of these receptors in AKH cells, caused changes in starvation survival and to locomotor activity. We further validated these by directly testing for changes in AKH cell physiology using live-cell imaging of explanted AKH cells. We found that these hormones evoked specific changes in fluorescent reporters in AKH cells. These results not only implicate novel functions for some of these hormones but also provide a better understanding of how multiple signals converge on AKH cells to modulate changes in organismal physiology.

## RESULTS

### AKH cells express multiple receptor molecules

To gain insight into potential regulatory elements governing AKH release, we evaluated the transcriptome of AKH cells. Individual AKH cells were identified by introducing a GFP reporter prior to microdissection and RNA sequencing. The transcriptome was assessed from 5 replicate individuals each that had experienced 24 hours of starvation and compared to animals that were fed *ad libitum*. An initial analysis of the RNA sequencing of 13 billion nucleotides corresponding to 11,456 transcripts showed no significant effects of the starvation experiments on the normalized levels of G protein-coupled receptor encoding genes, and consequently all replicates were pooled for further analysis.

We mined the AKH cell specific transcriptome to identify which cohort of neuropeptide and small molecule receptors were expressed specifically in AKH cells, to offer insight into potential endocrine connections that regulate AKH cell physiology and considered these two classes of receptor molecules separately. Given that the expression profile of the entire complement of neuropeptide receptor molecules was unchanged by nutrient conditions, we pooled the sequence reads across treatment to determine relative abundance of receptor expression. We found strong expression (as defined as counts in the top third of all genes) for 13 neuropeptide receptor genes, and consistent (as defined as counts in the top two thirds of all genes) expression for an additional 12 receptor genes (**Figure 1A**). Unsurprisingly, the most abundant neuropeptide receptor in AKH cells is the AstA-R1 receptor (**Figure 1A**), which had previously been reported to be an upstream regulator of AKH release (Hentze et al., 2015). Likewise, previous publications had implicated that the peptide Myosuppressin as an inhibitor of AKH secretion in different insects (Vullings et al., 1998), as well as the sNPF receptor (Oh et al., 2019). Thus, our analysis of the transcriptome shows consistency with previous behavioral results. Therefore, we focused on the relative high expression levels of the receptor that is specifically activated by the circadian hormone, PDF, as connections between the circadian system and metabolism are recognized in multiple taxa, however a connection between these two hormones have not yet been described.

**FIGURE 1.**
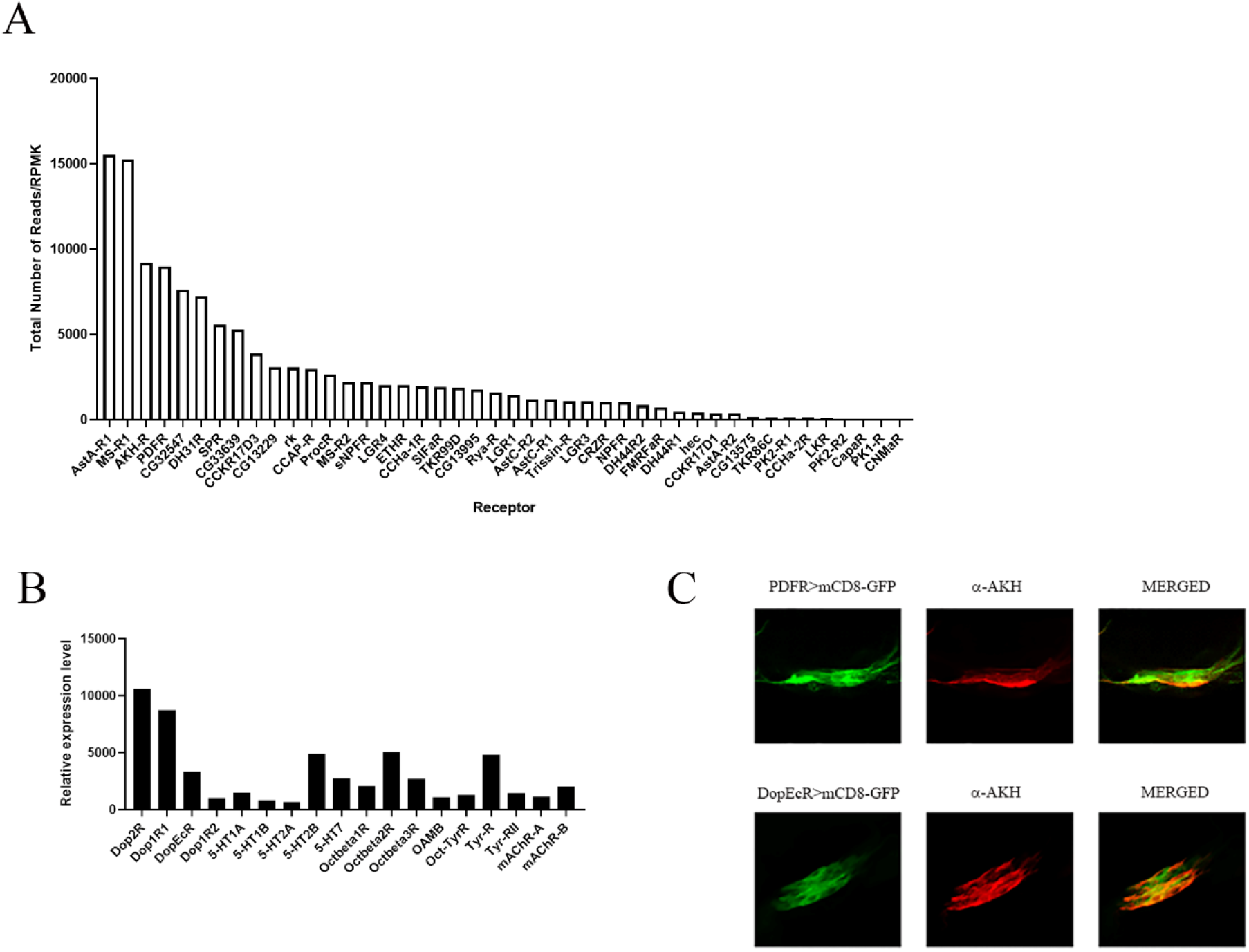
AKH cells express multiple GPCRs. Results from the RNAsequencing of individual APCs were mined for specific expression of neuropeptide GPCRs (1A) and amine GPCRs (1B). Each gene listed was reliably expressed across samples and the reads per million mapped reads are shown here for the expressed genes. Note that the PDFR is the 4th most abundant receptor expressed in APCs, following the Ast-A receptor, the MS-receptor and the AKH receptor. In 1B we note that APCs express multiple receptors for most of the small molecule transmitters, including octopamine, dopamine and serotonin. We validated expression for the PDFR in APCs by employing a specific GAL4 element that had previously been shown to rescue PDFR-phenotypes to drive GFP and using an antibody against AKH – show co-labeling (1C top). We used a similar approach for validating the DopEcR, using a DopEcR-Gal4 element to drive GFP and counterstained with AKH antisera. (1C. bottom).

In a set of complementary analyses, we assessed the specific expression of G protein coupled receptors that specifically bind small molecule transmitters. Interestingly, we found fairly uniform expression for nearly all the classical neurotransmitters (5HT, Octopamine, Dopamine, Acetylcholine, Tyramine, and Glutamate) (**Figure 1B**). AKH cells are known to modulate octopaminergic cells and vice versa (Yang et al., 2015, Pauls et al., 2020), and so our transcriptome verified previous experiments on AKH cell modulation. We chose to focus on dopamine receptor expression to test for potential modulation of AKH cell physiology, as many of the phenotypes associated with dopamine are similar to the starvation phenotypes that lie downstream of AKH signaling.

We first wanted to independently confirm that these molecules are expressed in AKH cells and employed two different methods. First, we employed single-cell RT PCR on dissected AKH cells and were able to specifically detect amplicons that corresponded to PDFR. Additionally, a PDFR driver element that has been shown to rescue relevant PDFR phenotypes (Lear et al., 2009) was used to introduce a GFP reporter to PDFR-expressing tissues. We used an AKH specific antibody (Braco et al. 2012) and found colocalization of the GFP reporter with the immunolabels (**Figure 1C**). Likewise, the DopEcR-GAL4 also showed strong expression in AKH cells.

### PDFR function in AKH cells is required for normal behavioral starvation responses

After validating that AKH cells express the PDFR, we next asked whether altered PDFR expression would lead to changes in starvation sensitivity, as altered physiology of AKH cells causes a variety of starvation phenotypes (Lee et al., 2004; Isabel et al., 2004). Multiple investigations consistently have found that the loss of AKH signaling leads to increased energy stores and decreased locomotion under starvation which together result in increased survival under starvation (Lee *et al.* 2004; Isabel *et al.* 2004; Braco *et al.* 2012). We hypothesized that if the PDF receptor was functioning to modulate the release of AKH, then the loss of the receptor would lead to a change in lifespan under starvation. Using this experimental framework, we tested the AKH cell specific introduction of an RNAi element targeting PDFR significantly for altered starvation behaviors. We found that PDFR knockdown lengthened starvation lifespan (**Figure 2A**), suggesting that PDFR facilitates AKH release. One interesting feature of PDFR is that it does not show desensitization in response to ligand binding and thus, persistent presentation of PDF would lead to chronic activation (Shafer *et al.* 2008; Choi *et al.* 2009). A genetically encoded membrane-tethered PDF (tPDF) has been useful to investigate PDF gain-of-function phenotypes and does so in only cells that endogenously express the PDFR (Choi *et al.* 2009a). We found that expression of tPDF in AKH cells resulted in a short-lived phenotype (**Figure 2A**). Since PDF is a known regulator of locomotor rhythmicity (Renn et al., 1999), and that AKH cells are critical for starvation-induced hyperactivity (Lee et al., 2004; Isabel et al., 2004), we asked if there were any changes in the locomotor profiles of these manipulations. PDFR-RNAi knockdown showed slightly elevated activity under replete conditions and a wild type response to starvation (**Figure 2B**). In contrast, expression of tPDF resulted in an abnormal locomotor pattern under both fed and starved conditions with especially high relative amounts of nighttime activity (**Figure 2B**). These results support the hypothesis that PDF facilitates AKH release, we next directly tested that hypothesis by assessing functional PDF signaling in AKH cells. The PDFR has been shown to signal predominantly through the cAMP second messenger system (Mertens et al., 2005), and endogenous PDFR activation has been visualized using the epac-camps reporter (Shafer et al., 2009). Therefore, we introduced this genetically encoded cAMP reporter to AKH cells and assessed whether AKH cells were responsive to the neuropeptide PDF. We found that direct application of PDF to explanted AKH cells decreased FRET signatures of the reporter, consistent with elevated levels of cAMP (Shafer *et al.* 2008). These values were similar to the FRET changes observed with application of forskolin, a positive control that elevates cAMP levels, and notably, there was a dose dependence of PDF on changes in FRET levels (**Figure 2C**). In order to test the directness of this response, we repeated these experiments with a co-expressed PDFR-RNAi element. Knockdown of PDFR in AKH cells abolished PDF-induced responses, showing that PDF is directly acting on AKH cells, and is dependent on the specific expression of the PDF receptor.

**Figure 2.**
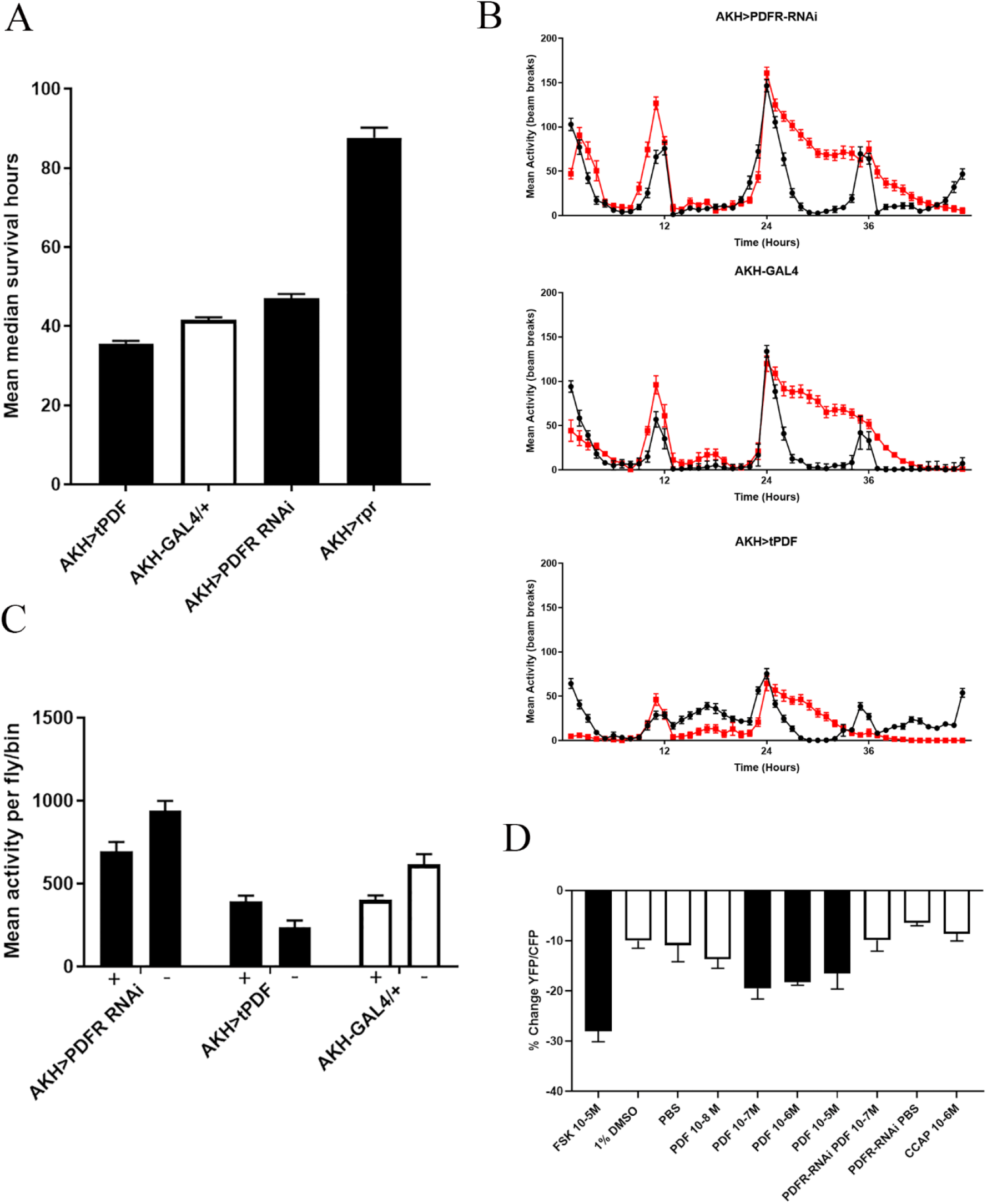
Manipulations of PDFR result in alterations of AKH dependent phenotypes. We genetically manipulated PDFR function in APCs and evaluated AKH related phenotypes including starvation lifespan (2A) and locomotor activity (2B and C). Figure 2A). Specifically, APC introduction of a PDFR-RNAi element lead to significantly longer mean lifespan during starvation) while introduction of a membrane-tethered PDF (t-PDF) to elicit constitutive PDFR signaling produced the opposite phenotype, specifically a significant shorter mean lifespan during starvation. (Black bars denote significant difference from controls (P< 0.05 ANOVA)). Figure 2B. Locomotor plots of locomotor activity during replete (black line) and starved (red line) conditions from animals expressing a PDFR-RNAi element (top) or a t-PDF element (bottom) as compared to genetic controls (middle). Figure 2C. Comparisons of total locomotor activity across these genotypes, Black bars denote statistical significance from control lines (P < 0.05 ANOVA). + denotes locomotor activity under replete conditions, and the – denotes activity during starvation. Figure 2D. Exogenous application of PDF alters the FRET signature of the epac-camps cAMP reporter. AKH cells expressing the epac-camps FRET sensor were isolated and different concentrations of PDF were applied. PDF elicited a significant change in FRET, consistent with previous demonstrations of this receptor modulating cAMP levels, and shows dose dependence. Notably, co-expression of a PDFR-RNAi element eliminates these responses. Black bars denote significant differences from vehicle addition.

### Dopamine receptor knockdown in AKH cells exhibit starvation phenotypes

Dopamine is a biogenic amine that mediates a number of different behaviors and physiologies in *Drosophila* (Friggi-Grelin et al., 2004). We thought it was an interesting observation that all known dopamine receptors were expressed in AKH cells and given some of the parallels in phenotypes, we tested for specific AKH phenotypes (starvation and locomotor activity) in animals expressing RNAi elements specifically targeting the different dopamine receptors. We found that genetic knock down of all four dopamine receptors in AKHs cells showed a significant reduction in lifespan under starvation (**Figure 3A**). Changes in lifespan were accompanied by aberrant locomotor phenotypes under starvation. We found that manipulations in three of the four dopamine receptors (DopEcR, Dop2R, and DopR) resulted in advanced onset of hyperactivity under starvation and overall increased locomotion, whereas conversely, D2R showed a significant decrease in locomotor activity (**Figure 3B**). Together these results suggest that dopamine may modulate AKH secretion, however, our results implicate multiple dopamine receptors involvement. It is also notable that one of these dopamine receptors, DopEcR has been shown to be a receptor for the steroid hormone, ecdysone, in addition to dopamine (Srivastava *et al.* 2005). To more fully explore the potential interaction of dopamine and ecdysone in AKH cells and to gauge the contributions of multiple dopamine receptors, we monitored explanted AKH cells expressing a calcium sensitive GFP, GCaMP6s1, in response to hormone application.

**Figure 3.**
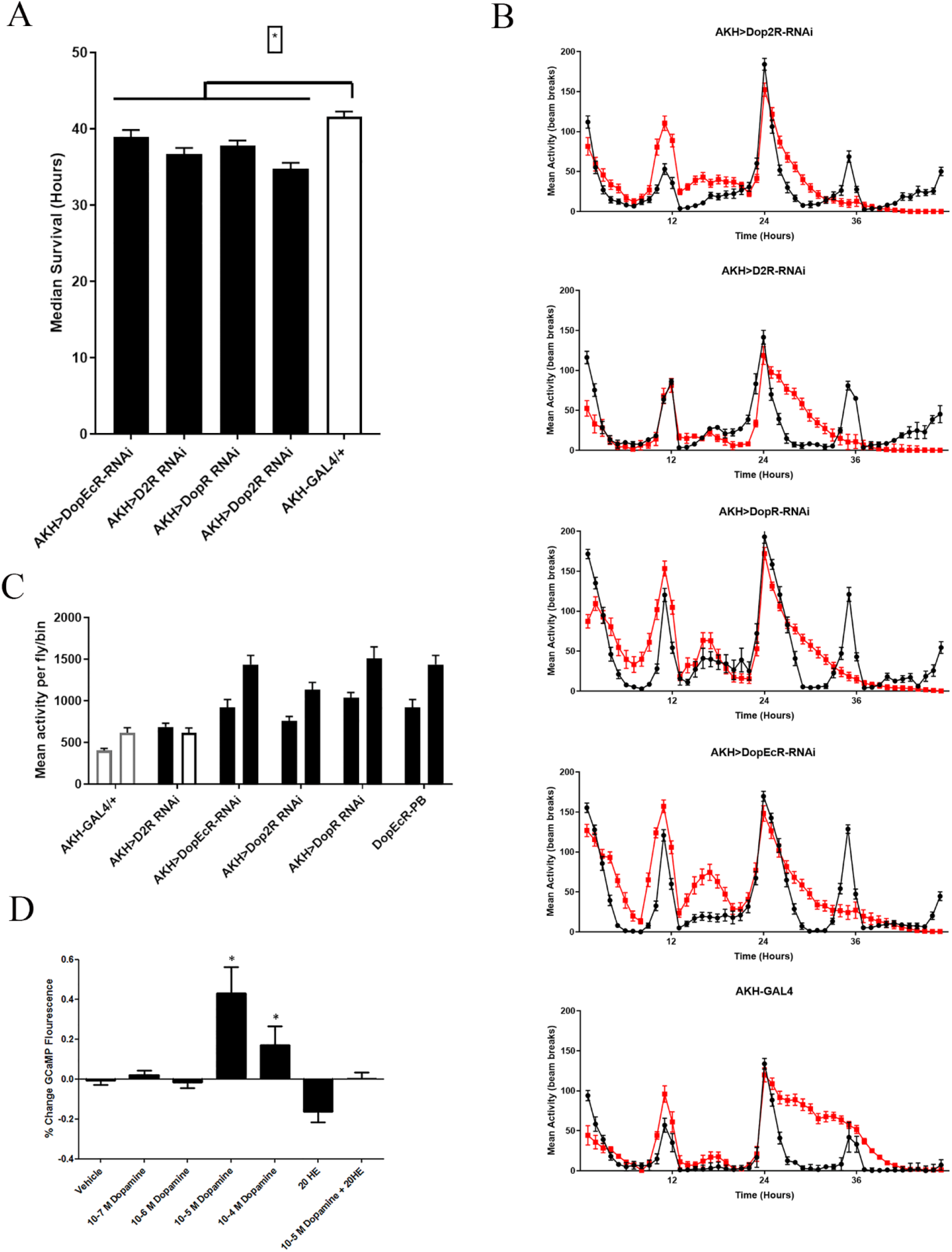
Dopamine modulates AKH dependent phenotypes. We introduced RNAi elements targeting each of the dopamine receptor present in AKH cells and assessed starvation lifespan (Figure 3A) and locomotor activity (Figure 3B and 3C). We note that the each of the dopamine receptor RNAi elements significantly reduced lifespan during starvation (black bars denote significance P <0.05, ANOVA). Figure 3B. Locomotor plots of locomotor activity during replete (black line) and starved (red line) conditions from animals expressing RNAi elements targeting a specified Dopamine receptor as compared to genetic controls. Figure 2C. Comparisons of total locomotor activity across these genotypes, Black bars denote statistical significance from control lines (P < 0.05 ANOVA). + denotes locomotor activity under replete conditions, and the – denotes activity during starvation. Figure 2D. Exogenous application of dopamine alters GCaMP reporter fluorescence. AKH cells expressing the GCaMP sensor were isolated and different concentrations of DA were applied. DA elicited a significant change in GCaMP and showed dose dependence. Notably, co-application of ecdysone eliminates these responses. Black bars denote significant differences from vehicle addition.

We next tested if dopamine was capable of effecting calcium levels in explanted AKH cells. Interestingly, we found that dopamine-induced responses were dependent on the extracellular concentration of sugar. Under high extracellular trehalose levels (15mM), dopamine application failed to produce a response, however under low extracellular trehalose levels (3mM), dopamine induced a strong peak in calcium in a dose-dependent manner (**Figure 3C**). We hypothesize that this context dependence likely reflects changes in basal receptivity of AKH cells. We, and others, have previously reported that low extracellular trehalose concentration result in AKH cell activation, however this was over a much longer time course (30 minutes), whereas peak dopamine responses occurred within 30 seconds after application. We next tested if ecdysone was capable of changing calcium concentrations and found that application of 20HE (20-hydroxyecdysone) resulted in no change in AKH cell activation. However, co-application of ecdysone and dopamine completely blocked any increase in calcium suggesting that these two hormones are acting antagonistically *in vivo* (**Figure 3C**).

## DISCUSSION

In this study, we identified and characterized three hormones, (dopamine, ecdysone, and PDF) that are likely to directly regulate AKH secretion. Our initial exploration of the AKH cell transcriptome revealed the expression of a number of potential candidate GPCRs. We validated the expression of these receptors using multiple genetic and molecular methods. Next, we tested the behavioral consequences of knocking down these receptors in AKH cells and found that knockdown of each dopamine receptor produced aberrant changes in locomotion. Specifically, we found that three of the four dopamine receptors showed increase baseline locomotion as well as earlier onset of starvation induced hyperactivity. These manipulations also manifested in short lived phenotypes under starvation. We also found that changes in PDF signaling resulted in changes in both lifespan and locomotion during nutrient deprivation. These results were further supported by a functionally characterization of dopamine- and PDF-evoked responses in AKH cells using live-cell imaging. Collectively, these results clearly show that AKH cells are a critical point of integration lying at the crossroads between physiology/behavior and metabolic state of the organism.

Our experiments identified Pigment Dispersing Factor, PDF, as a candidate hormone that regulates AKH cell physiology. PDF is best known for its paramount role in regulating circadian rhythmicity (Renn et al., 2001; Hyun *et al.* 2005; Shafer *et al.* 2008; Choi *et al.* 2009b; Peschel and Helfrich-Förster 2011). We postulated that PDF may be acting to regulate AKH cells in a circadian fashion, as recent studies have established connections between the circadian system and metabolic control (Sehgal 2016). The current model of PDF action is that of a wake-promoting hormone, secreted in the early morning, and peaking in concentration before dawn (Schneider and Stengl 2005). Based on the evidence of the PDF receptor (PDFR) being expressed in AKH cells, we speculated that PDF may target AKH cells to coordinate energy release in anticipation of morning activity. Introduction of an RNAi element targeting PDFR significantly extended lifespan under starvation and in contrast, the introduction of the tethered PDF (to constitutively stimulate PDFR) decreased starvation longevity. These observations are consistent with a model in which PDF facilitates AKH release.

Since PDF is a responsible for the temporal regulation of locomotor activity, we next analyzed the locomotor activity under both replete and starvation conditions. Surprisingly, we found PDFR knockdown showed elevated locomotion under replete conditions and in contrast, expression of tPDF showed significantly lower basal activity that appeared unchanged during starvation. Considering that AKH is a requirement for starvation-induced hyperactivity and that it appears that AKH gain of function variants enhance locomotor activity (Isabel *et al.* 2005), these results appear to contradict the interpretation that PDF acts to increase AKH titers. Notably, animals with constitutive PDFR activation showed abnormally high levels of activity during the night and perhaps, the temporal change in activity levels reflects a chronic sleep deprivation, and this is the explanation for reduced longevity. Notably, we observed that exogenous PDF application to explanted AKH cells result in increased levels of cAMP, the second messenger downstream of PDFR and that such responses were PDFR dependent. Collectively, these results verify that PDFR is expressed in AKH cells and that PDFR action regulates AKH release.

Our work, as well as previous reports, suggest that AKH is dispensable for rhythmic locomotor response (Lee *et al.* 2004). Although, we cannot fully rule out the potential for PDF to modulate AKH in a circadian manner, the absence of any circadian phenotype in locomotion suggests that PDF may be acting on AKH cells in a clock-independent fashion. A major hypothesis of PDF action is that it acts to synchronize other temporal centers (Peschel and Helfrich-Förster 2011; Seluzicki *et al.* 2014). However, there are groups of non-clock cells that regulate rhythmic feeding *via* PDF modulation (Cavey *et al.* 2016). and other investigations suggest that PDF may have non-clock functions. Specifically, manipulations of PDF signaling alters triglyceride levels independent of clock function (DiAngelo *et al.* 2011). Given that the circadian loci of PDF action has been mapped to a group of central neurons (Shafer *et al.* 2009), it seems unlikely that central PDF neurons are impacting AKH cells. In addition to central expression, PDF is also expressed in a subset of neurons in the ventral nerve cord (VNC) that project posteriorly to the midgut (Talsma *et al.* 2012). These PDF neurons do not express any clock genes and are dispensable for circadian rhythmicity. Furthermore, these same neurons have been shown to impact ureter contractions in the midgut and osmotic homeostasis (Talsma *et al.* 2012). It is possible that these PDF neurons may provide the hormone source that is upstream of AKH. PDFR is also known to respond to another ligand DH31, although comparatively with lower affinity (Johnson *et al.* 2005; Mertens *et al.* 2005). DH31 is a multifunctional hormone that has been linked to stress response behaviors as well as circadian rhythms (Kunst *et al.* 2014). Unlike PDF, DH31 is expressed in many tissues, including enteroendocrine cells in the gut (Park *et al.* 2011) and therefore, may be more abundant in circulating hemolymph than PDF. Consequently, it may be that DH31 signaling via PDFR that is biologically relevant to AKH cells. While the precise mechanism for PDFR activation in AKH cells remains unclear and likely to be the subject of future studies, our results firmly cement that PDFR modulates this metabolic center.

Previous literature has found that dopamine is involved in a number of different behaviors and physiologies including courtship, memory formation, and circadian rhythms (Lebestky *et al.* 2009; Keleman *et al.* 2012; Linford *et al.* 2015b). Interestingly, functional characterization of dopamine signaling illustrates numerous parallels with AKH signaling. Specifically, pharmacological and genetic manipulations, intended to elevate dopamine signaling, cause increased locomotor activity and enhanced gustatory perception of sugar (Kume *et al.* 2005; Inagaki *et al.* 2012; Yamamoto and Seto 2014). Dopamine levels also rise in response to multiple forms of stress including oxidative, heat shock, and starvation (Gruntenko and Rauschenbach 2008).

These experiments also implicate ecdysone signaling as an important endocrine factor that regulates metabolic status. Either a reduction of the expression of DopEcR levels or the complete loss of the receptor both show changes in locomotion levels and starvation sensitivity. The DopEcR is an interesting GPCR as it also binds the lipid-soluble steroid, ecdysone (Srivastava *et al.* 2005). Specifically, ecdysone has been shown to abolish dopamine-induced responses (Srivastava *et al.* 2005) and is thought to change the signaling parameters of the receptor (Evans *et al.* 2014). While ecdysone signaling is well-known for its impact in mediating developmental transitions, studies have shown ecdysone signaling to be a critical mediator of stress responses in adult insects (Simon *et al.* 2003; Terashima *et al.* 2005).

Like dopamine and AKH, ecdysone titers increase under metabolic stress (Gruntenko and Rauschenbach 2008). Furthermore, increased ecdysone halts oogenesis in females and consequently, shifts nutrient allocation from reproduction to survival (Terashima *et al.* 2005). From this evidence we hypothesized that ecdysone may abolish the inhibitory effects of dopamine on AKH cells and directly tested for interactions of these two hormones. We found that administration of dopamine under high extracellular trehalose failed to induce any change in GCaMP fluorescence. However, we did find that when extracellular trehalose was low, dopamine induced a strong increase in fluorescence. Furthermore, co-application of 20E eliminated any dopamine response under these nutrient levels. Ecdysone is thought to act by directly modulating DopEcR activity, in our experiment the entirety of the dopamine-induced response was abolished by ecdysone even though other dopamine receptors are present in AKH cells. One potential hypothesis is that the other dopamine receptors signal through different mechanisms so that the calcium influx is primarily downstream of the DopEcR. Consistent with that hypothesis, the DopEcR is the most abundantly expressed dopamine receptor according to our transcriptome dataset. Also it has been previously suggested that ecdysone doesn’t inhibit dopamine signaling *per se*, but rather changes the second messenger that is downstream of receptor activation (Evans et al., 2014) and our observations would be consistent with that model of DopEcR activation. One other potential mechanism that can explain these results, is that the other dopamine receptors may act to modulate DopEcR signaling to precisely regulate AKH cell responsiveness to these different ligands (dopamine or ecdysone). Future experiments aimed at identifying the roles of these molecules are required, as our results clearly demonstrate that dopamine and ecdysone are acting antagonistically on AKH cells.

While all of our independent experiments firmly establish a role for dopamine and ecdysone impacting AKH signaling, how do we reconcile the different results suggesting that dopamine may inhibit or enhance AKH signaling? One hypothesis is that dopamine may act as a neuromodulator, and the idea that dopamine must either be an excitatory or an inhibitory input is likely too simplistic. That idea is supported by the observation that multiple dopamine receptor subtypes are present in AKH cells, and that dopamine responsiveness of AKH cells is dependent on extracellular sugar levels. Furthermore, interpretation of our behavioral experiments center on animals expressing lifelong genetic constructs which may not be readily comparable to interpretations of dopamine action on AKH cell physiology observed along much shorter timescales. Given the contextual dependence of dopamine signaling, we submit that loss of dopamine receptor signaling phenotypes may not simply distill into a model in which dopamine inhibits AKH secretion. Further complicating our understanding of the behavioral phenotypes is the fact that both dopamine and ecdysone rise in response to stress *in vivo*. Our results suggest that these two hormones are acting antagonistically to regulate AKH indicating that the pertinent information is in the ratio of these two hormones. Consequently, the net contribution of this receptor to AKH cells is reliant on the precise stoichiometry of these two molecules at a given time. Furthermore, the temporal dynamics of both dopamine and ecdysone increases are unclear and it may be that the relevant information is when during stress these hormones are released as well as absolute abundance.

These results demonstrate that AKH cells are regulated by multiple hormones. While the predicted phenotypic outcome proved more complex that initially hypothesized it is clear that dopamine, ecdysone, and PDF are all capable of directly acting on AKH cells. Furthermore, our results suggest that these signals are interacting in a complex context-specific manner. This study serves as an experimental foundation to further unravel the mechanisms underlying AKH cell physiology and the endocrine circuits that modulate metabolism and maintain energetic homeostasis.

## MATERIAL AND METHODS

### Fly husbandry

All flies were maintained in an incubator maintained at 25° and under a 12:12 light/dark (LD) cycle unless otherwise stated. Flies were cultured on a standard molasses–malt–cornmeal– agar–yeast medium and housed in uncrowded conditions (Zhao et al., 2010). We used the following fly strains: PDF-GAL4 (Renn et al., 2004)(BL-6869), AKH-GAL4 (Lee et al., 2004) (BL-25684),UAS-tPDF (Choi et al.,2009), PDFR-GAL4 (Lear et al., 2004) (BL-30370), UAS-epac-camps (Shafer et al., 2008)(BL-25407), UAS-mCD8-GFP (BL-5137), UAS-PDFR-RNAi (BL-38347), UAS-DopEcR-RNAi (BL-31981), UAS-Dop2R-RNAi (BL-51423), UAS-D2R-RNAi (BL-50621), UAS-DopR-RNAi (BL-55239), UAS-GCaMP6s1 (BL-42746).

### AKH cell specific Transcriptome

AKH cells expressing GFP under the AKH promoter were microdissected and aspirated into a glass pipette, which was placed in a PCR tube and flash frozen in an ethanol-dry ice bath. The tubes were stored at −80 for no longer than three weeks while 10 samples were prepared from 5 fed and 5 starved flies. On the day of RNA amplification, the contents of the PCR tubes were centrifuged and the RNA from these samples was amplified in parallel with the Arcturus RiboAmp HS PLUS Kit by following the manufacturers protocol (KIT0505, Thermo Fisher Scientific). RNA libraries were then prepared using the Kapa Stranded mRNA-Seq library prep kit and 50 bp single end sequencing was performed on an Illumina HiSeq 4000 at Duke Center for Genomic and Computational Biology (Durham, NC). This data is available at the NCBI Sequence Read Archive under project number PRJNA642982.

The raw reads were filtered using Trimmomatic v0.36 to remove Illumina adaptors, leading or trailing bases below a quality score of 3, 4-base sliding window average quality below 15 and reads less than 36 bp long. Filtered reads were aligned to *Drosophila melanogaster* genome BDGP6.22 using star v2.5 (Dobin et al., 2013) and a count table generated from coordinate sorted BAM files using summarizeOverlaps from the biocondutor package GenomicAlignments (Lawrence et al., 2013). We identified genes differentially expressed by starvation using the Bioconductor package DESeq2 (Love et al., 2014), but no GPCRs were differentially expressed under starvation conditions (adjusted p > 0.05). The source code for this analysis is available at github (Saunders, 2020).

### Individual locomotion/starvation

Three to five-day old males were sorted 12-24 hours prior to the start of the assay. At ZT0 individual male flies were loaded into 5 x 65 mm Polycarbonate Plastic tubes capped at one end with a ½ inch piece of yarn. Once loaded, a 200 μL pipette tip filled with standard *Drosophila* media and sealed at one end was placed on the end of the plastic tube. Tubes were then loaded into Trikentics DAM 2 monitor for 3 days of entrainment on replete media. Total beam counts were monitored continuously through an automated system for the duration of the experiment at 10-minute intervals. At ZT0 on the third day data collection was paused and media containing pipette tips were replaced with a tip containing a 2% agar water solution. Locomotion was monitored for at least three days or until all flies in starvation had ceased moving for 12 hours. Following the experiment, beam breaks were binned into 1 hour intervals and used for locomotor analysis. Day 1 data was considered a recovery and acclimation period and was removed from analysis. Death was approximated as the time point following the last registered beam break.

### Live cell imaging

For live cell imaging experiments, adult ring glands were dissected and placed in AHL (adult hemolymph-like) (Feng *et al.*, 2004) solution containing 12mM trehalose and 3Mm sucrose or 3mM trehalose 12mM sucrose (when stated). Dissections where then placed on a plastic cover slip containing 180 μL of AHL. Explanted ring glands were then viewed on a Zeiss LSM 710 confocal microscope and visually inspected for damage prior to imaging. All imaging settings were kept constant between experiments.

For calcium imaging a 20x 0.8 NA objective and a 488nm laser were used. Z stacks were collected in 10 second intervals. Cells were imaged for 1 minute prior to treatment. After imaging Z stacks were collapsed to maximum intensity projections. A region of interest was manually drawn for each ring gland and total values for pixel intensity were assessed. Values were exported in Excel and normalized to the time point immediately prior to application. Dopamine was prepared in 10% PBS AHL solution. A Kruskal Wallis ANOVA was used for analysis and no fewer than 5 replicates were tested per condition.

For cAMP imaging, a 40X 0.95NA objective was used. CFP was excited using a 440nm laser. Z stacks were collected in 10 second intervals and imaged for 1 minute prior to treatment. After imaging, Z stacks were collapsed to maximum intensity projections. A region of interest was manually drawn for each ring gland and total values for pixel intensity were assessed. Values were exported in Excel and adjusted for spillover (SO). Spillover was calculated using CFP expressing HEK cells under the same conditions previously described and found to be 54%. Fret ratio was calculated 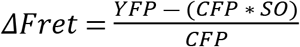 (Shafer *et al.* 2008). Data was normalized to the time point immediately prior to application and Friedman’s test was used to determine significance. No fewer than 3 replicates per condition were tested.

### Immunostaining

All tissues dissections for immunostaining were done in 1X PBS Tx under a standard dissecting microscope. Tissue was then placed immediately into fixative (4% Paraformaldehyde 7% picric acid) for 1 hour at room temperature. Tissue was then washed 10X with 1X PBS Tx before blocking with (%) BSA 1X PBS Tx solution for one hour at room temperature. Next tissue was placed in 1:1000 αAKH for 1 hour at room temperature and then washed 10X before moving the tissue to 1:1000 Anti Rabbit Cy3 for 2 hours at room temperature. Tissue was next washed and placed into a drop of glycerol to dehydrate it. Finally, the glycerol was removed by a kimiwipe (wicking) and replaced with Anti Fade mounting media. All images were taken on a Zeiss 710 confocal microscope.

## Acknowledgements

This work was supported by NSF IOS1355097 and a Center for Molecular Signaling grant to ECJ. We thank the Bloomington Stock Center NS Dr. Michael Nitabach for fly stocks and acknowledge the Distributed Environment for Academic Computing (DEAC) at Wake Forest University for providing HPC resources that have contributed to the research results reported within this paper. URL: https://is.wfu.edu/deac

## AUTHOR STATEMENT

The precise mechanisms of energy allocation in organisms is not completely understood. In Drosophila, the Adipokinetic Hormone which is the functional equivalent of mammalian glucagon features prominently in mediating behavioral and physiological changes accompanying nutrient limitation. In this study, we examine the AKH cell specific transcriptome and find multiple receptor molecules expressed in this cell type. We find that the circadian hormone, Pigment Dispersing Factor and the biogenic amine, dopamine, alter AKH cell physiology and suggest a complex endocrine circuit that modulate basic metabolism.

